# The neural basis of precise visual short-term memory for complex recognisable objects

**DOI:** 10.1101/085068

**Authors:** Michele Veldsman, Daniel J. Mitchell, Rhodri Cusack

## Abstract

Recent evidence suggests that visual short-term memory (VSTM) capacity estimated using simple objects, such as colours and oriented bars, may not generalise well to more naturalistic stimuli. More visual detail can be stored in VSTM when complex, recognisable objects are maintained compared to simple objects. It is not yet known if it is recognisability that enhances memory precision, nor whether maintenance of recognisable objects is achieved with the same network of brain regions supporting maintenance of simple objects.

We used a novel stimulus generation method to parametrically warp photographic images along a continuum, allowing separate estimation of the precision of memory representations and the number of items retained. The stimulus generation method was also designed to create unrecognisable, though perceptually matched, stimuli, to investigate the impact of recognisability on VSTM. We adapted the widely-used change detection and continuous report paradigms for use with complex, photographic images.

Across three functional magnetic resonance imaging (fMRI) experiments, we demonstrated greater precision for recognisable objects in VSTM compared to unrecognisable objects. This clear behavioural advantage was not the result of recruitment of additional brain regions, or of stronger mean activity within the core network. Representational similarity analysis revealed greater variability across item repetitions in the representations of recognisable, compared to unrecognisable complex objects. We therefore propose that a richer range of neural representations support VSTM for complex recognisable objects.

## Introduction

Visual short-term memory (VSTM) enables us to operate in our rich and dynamic visual environment. In the last decade, there have been many advances in theoretical understanding of VSTM (Luck and Vogel 2013) but these have largely been based upon experiments using simple abstract stimuli such as coloured patches and geometric shapes that lack important facets of complex visual objects in the environment (Brady et al. 2011). Most saliently, objects in the environment are frequently recognizable, allowing rich information to be retrieved from long-term memory, such as their category, prototypical form, and semantics. But, does this influence our memory for their visual features?

Recently, Brady et al. (2016) demonstrated quantitatively different VSTM capacity for complex real world objects compared to simple colours. More complex items could be stored with finer detail than simple colours when encoding time was increased. But simple colours and complex objects differ in both semantic and perceptual complexity, therefore careful control of visual features is important in assessing the quality and capacity of VSTM for complex objects and the impact of recognisability. Similarly, it remains to be determined how complex, recognisable objects in VSTM are represented in the brain compared to perceptually matched, complex, but unrecognisable items.

Brady et al. (2016) showed that maintenance of complex items was associated with higher amplitude contralateral delay activity (CDA), an electrophysiological marker that indexes the amount of visual information actively stored. Contrary to the expectation that episodic long-term memory would support the maintenance of complex recognisable objects, Brady et al. (2016) argued that the same mechanism supporting maintenance of simple objects, indexed by the CDA, also supported maintenance of complex objects. This raises the question as to whether the same spatial network of regions recruited in the maintenance of simple objects is also sufficient to support maintenance of complex recognisable objects when they are perceptually matched. A meta-analysis of change detection tasks showed activity across a network of frontal and parietal regions in the encoding and maintenance of simple objects (Linke et al. 2011). VSTM for complex recognisable objects, with their semantic associations, may recruit the same regions to a greater extent or recruit additional regions, such as those associated with long term memory. Alternatively, recognisability may modulate fine-scale activity patterns within the core network.

Our first goal was to determine how VSTM for complex objects is affected by how recognizable they are. To study this, we created sets of variably recognizable stimuli that were matched in their visual properties. We used a warping procedure that made photographs of objects more difficult to recognize, but across the set gave indistinguishable distributions of responses in computational models of early visual processing (Stojanoski & Cusack, 2014).

Our second goal was to understand the neural mechanism through which recognisability affects VSTM. We hypothesized that visual regions may be tuned for recognizable objects, which may allow for more efficient neural codes that are more effectively maintained in VSTM. A second hypothesis was that the recognition of an object could recruit additional regions in the hierarchically organised ventral visual stream (DiCarlo et al. 2012). These additional regions might directly contribute to VSTM, or might indirectly support representations in earlier visual regions.

To preview the results, across three experiments (total N=88) we demonstrate that the visual features of recognisable objects are remembered with higher precision than those of unrecognisable objects. However, we found no consistent evidence that this clear behavioural benefit was supported by increased regional brain activation, or recruitment of additional brain regions. Instead, multivariate pattern analysis revealed greater variability in the representation of recognisable objects, suggesting a wider representational space may support VSTM for complex, recognisable objects.

## 1 Experiment 1

In experiment 1 we measured the effect of recognisability on VSTM, and the neural basis of this effect. We first compared the detection of large (cross-category) versus small (within-category) changes. This manipulation has previously been applied a number of times to assess the precision of a VSTM representation from the likelihood of an item being remembered at all (Awh et al. 2007; Scolari et al. 2008; Barton et al. 2009).

### 1.1 Methods

#### 1.1.1 Participants

Eighteen participants (9 male, aged 18-46, mean age 28) gave informed consent, approved by the Cambridge Research Ethics Committee, and were paid for taking part. Participants had no history of psychological or neurological health problems and reported normal or corrected to normal vision.

#### 1.1.2 Stimuli

Forty colour photographic images of real world objects (21 depicting inanimate objects) were selected from the stimuli used in Kriegeskorte, Mur, Ruff, et al. (2008). Stimuli were then manipulated in Matlab (MathWorks, 2009B). A diffeomorphic transformation was applied to each photographic image to parametrically degrade its recognisability. Diffeomorphic warping was preferred over procedures such as phase, box or texture scrambling that are also designed to remove recognition, because it provides better matching of visual properties (Stojanoski and Cusack 2014). Like bending a rubber sheet, the diffeomorphic transformation maintains a 1:1 mapping between each point in the source and a point in the target, in a continuous transformation across space, without replacement or duplication. The transformation is also smooth and reversible (see Stojanoski and Cusack (2014) for full details of the transformation).

#### 1.1.3 Change Detection Paradigm

We used the common change detection paradigm to measure VSTM (Cowan 2001; Scolari et al. 2008). A single sample item was presented centrally for 0.25s spanning a visual angle of 3.70⁰ on a uniform grey background (Fig 1a). The sample was either intact (recognisable) or heavily distorted (unrecognisable). Memory load was fixed to a single item. The sample was followed by a jittered maintenance period (selected from a random distribution between 1-9 seconds) in which a uniform grey background was displayed. A probe was then presented centrally for 1.5s, which matched the sample item on half of the trials and changed on half of the trials. Participants made a ‘same’ or ‘different’ response with a response button box in the scanner. Response mapping was counterbalanced across participants. A blank inter-trial-interval followed the probe, whose duration was selected from a random distribution between 1-9 seconds. The jittered inter-trial and inter-stimulus periods were designed to allow separation of the encoding, maintenance and response periods (Rowe and Passingham 2001).

**Fig 1.**
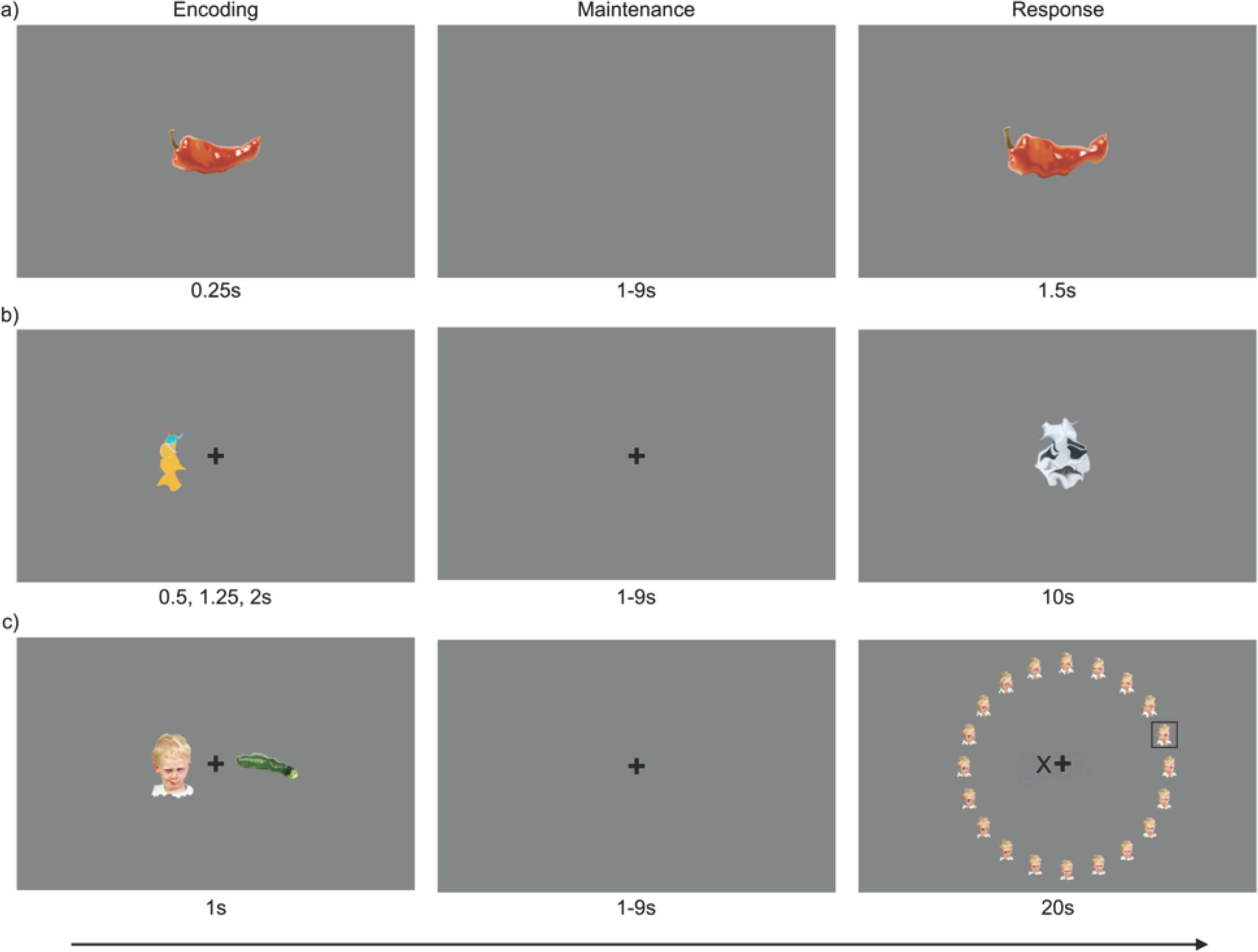
a) Experiment 1; a single stimulus is displayed (recognisable trial in example) followed by a jittered maintenance period (1-9s). Participants indicate whether the probed item is the ‘same’ or ‘different’ (within-category change in example) b) Experiment 2; one or two items displayed at encoding (single unrecognisable trial in example) followed by a jittered maintenance period and a centrally presented probe (cross-category change condition in example). c) Continuous report task. Participants used a selection box to choose from the response wheel the item most similar to the remembered item on the side cued with the X.

To distinguish between complete forgetting, imprecise remembering, and precise remembering, we manipulated the type of change to be detected (Scolari et al, 2008). In cross-category (CC) trials the identity of the stimulus changed between the sample and probe, with the level of recognisability (degree of diffeomorphic distortion) maintained. In within-category (WC) trials the level of recognisability changed between the sample and the probe, with the identity of the sample maintained. 40 CC and 40 WC trials were blocked and the order of blocks was counterbalanced across participants. Within each block, change/no-change trials were counterbalanced with the recognisability manipulation, and presented in a random order. VSTM capacity was estimated using Cowan’s formula (*hits-false alarms)/set size*; Rouder et al. 2011) which gives a measure of the items-worth of information retained (K).

The task was presented to participants with a PC running the Psychophysics Toolbox Version 3 (Brainard, 1997) extension of Matlab (Mathworks, Natick, MA). This was projected into the scanner using a Christie (Cypress, CA) video projector at a 60 Hz refresh rate and viewed in a mirror approximately 90 mm from the participants’ eyes.

#### 1.1.4 MRI Acquisition

Scanning took place at the Medical Research Council Cognition and Brain Sciences Unit, Cambridge on a 3T Siemens TIM Trio (Erlangen, Germany). For fMRI, T2*-weighted echo-planar images (EPI) were acquired (Repetition time (TR) 2s; echo time (TE) 30s; flip angle 78°; 32, 3.5mm slices with 10% gap, 64×64 acquisition matrix and 3 × 3 × 3.75mm voxel size). The first 10 seconds of scans were discarded to allow for T1 equilibrium. For anatomical localisation, a high-resolution T1-weighted 3D MPRAGE was acquired with TR of 2.25 secs; TE of 2.99ms; TI of 900ms, 9⁰ flip angle, 256 × 240 ×192 matrix size and 1mm isotropic voxels.

#### 1.1.5 Image processing and analysis

Automated processing software (aa,www.github.com/rhodricusack/automaticanalysis) was used to preprocess and analyse the functional imaging data (Cusack et al. 2014) in SPM8 (Wellcome Department of Imaging Neuroscience, London, UK; http://www.fil.ion.ucl.ac.uk/spm/). The preprocessing pipeline comprised motion correction, slice-time correction, coregistration to the individual’s structural image, normalisation to MNI template space and smoothing with a 10 mm FWHM Gaussian kernel. A high pass filter with a cut-off of 128 s was applied to remove low frequency noise. Regressors of interest modelled recognisable and unrecognisable target trials and CC and WC change trials in the encoding, maintenance and response periods of the task. Events were convolved with the canonical hemodynamic response function and a general linear model (GLM) was fitted voxel-wise. Six parameter motion estimates were included in the model to account for noise related to head motion in the scanner. Individual participant data were entered into a group level random effects analysis.

Whole-brain voxel-wise analyses were conducted and tested using the false discovery rate (FDR) correction for multiple comparisons at a threshold of α<0.05 unless otherwise stated. Results were visualised with BrainNet Viewer (Xia et al. 2013). Main effects contrasts in the whole brain and ROI analyses are relative to an implicit baseline, that is all remaining events not included in the model itself, essentially the inter-trial-intervals.

A region of interest (ROI) analysis was conducted to enable comparison of brain activity associated with VSTM for complex objects across the paradigms presented here and with the meta-analysis of VSTM imaging studies by Linke et al., (2011). Details of the regions used are reported in Linke et al., (2011). The ROIs include regions of the inferior intraparietal sulcus (inf-IPS) and superior intraparietal sulcus (sup-IPS, MD-IPS and Silver-IPS), inferior frontal sulcus (IFS), middle frontal gyrus (MFG) and lateral occipital cortex (LOC; see Fig 4 and 6 for visualisation) that are reliably activated in VSTM tasks of simple objects. ‘Silver-IPS’ and ‘MD-IPS’ are so called to distinguish them the inferior and superior IPS regions from Xu & Chun (2006). “Silver-IPS” refers to an ROI taken from Silver et al. (2005) which examined topographic maps of visual attention in parietal cortex. The “MD-IPS” refers to a node of the “multiple-demand” network, a collection of regions commonly activated across a wide variety of tasks, including VSTM tasks (Duncan and Owen 2000). ROIs were created as 10 mm spheres around peak MNI coordinates reported in Linke et al. (2011).

**Fig 2.**
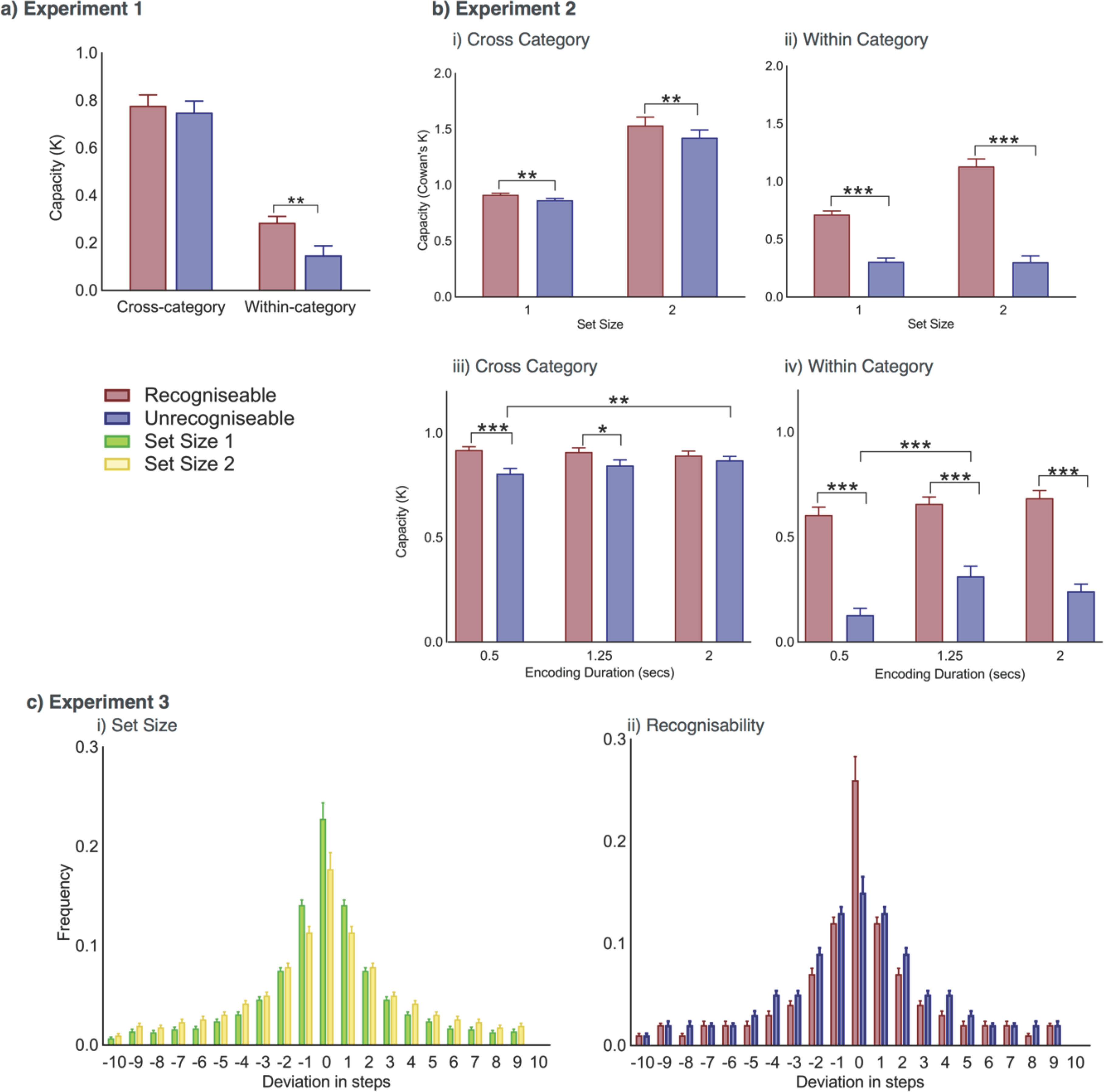
a) Mean capacity (K) in Experiment 1 for recognisable and unrecognisable items in the cross-category (CC) and within-category (WC) condition. b) Mean capacity (K) in Experiment 2 in the i) CC and ii) WC condition separated by recognisability, and in the iii) CC and iv) WC condition separated by encoding duration. c) Experiment 3; Distribution of responses in deviation in steps from the target response value i) for set size 1 and 2 trials and ii) recognisable and unrecognisable target item trials. Error bars indicate standard error. * p<.05; **p<.01; ***p<.001.

**Fig 3.**
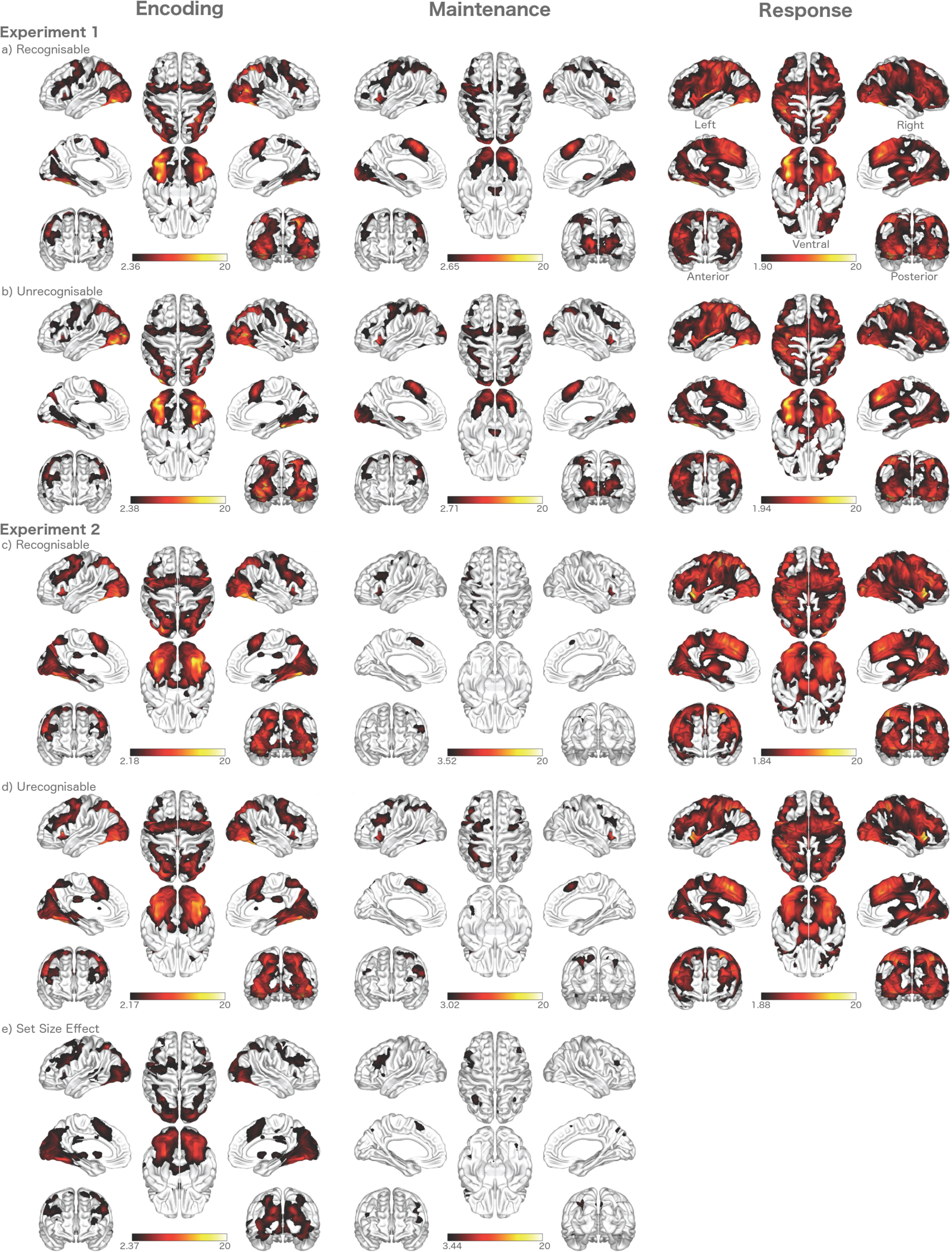
Experiment 1: Significant activity associated with recognisable object trials (row a) and unrecognisable object trials (row b) in the encoding, maintenance and response phases. Experiment 2: Significant activity associated with recognisable trials (row c) and unrecognisable trials (row d) in the encoding, maintenance and response phase. Significantly greater activity in the encoding and maintenance of two items compared to a single item (row e). Colour bars represent t-values. All contrasts are relative to implicit baseline. FDR <0.05.

**Fig 4.**
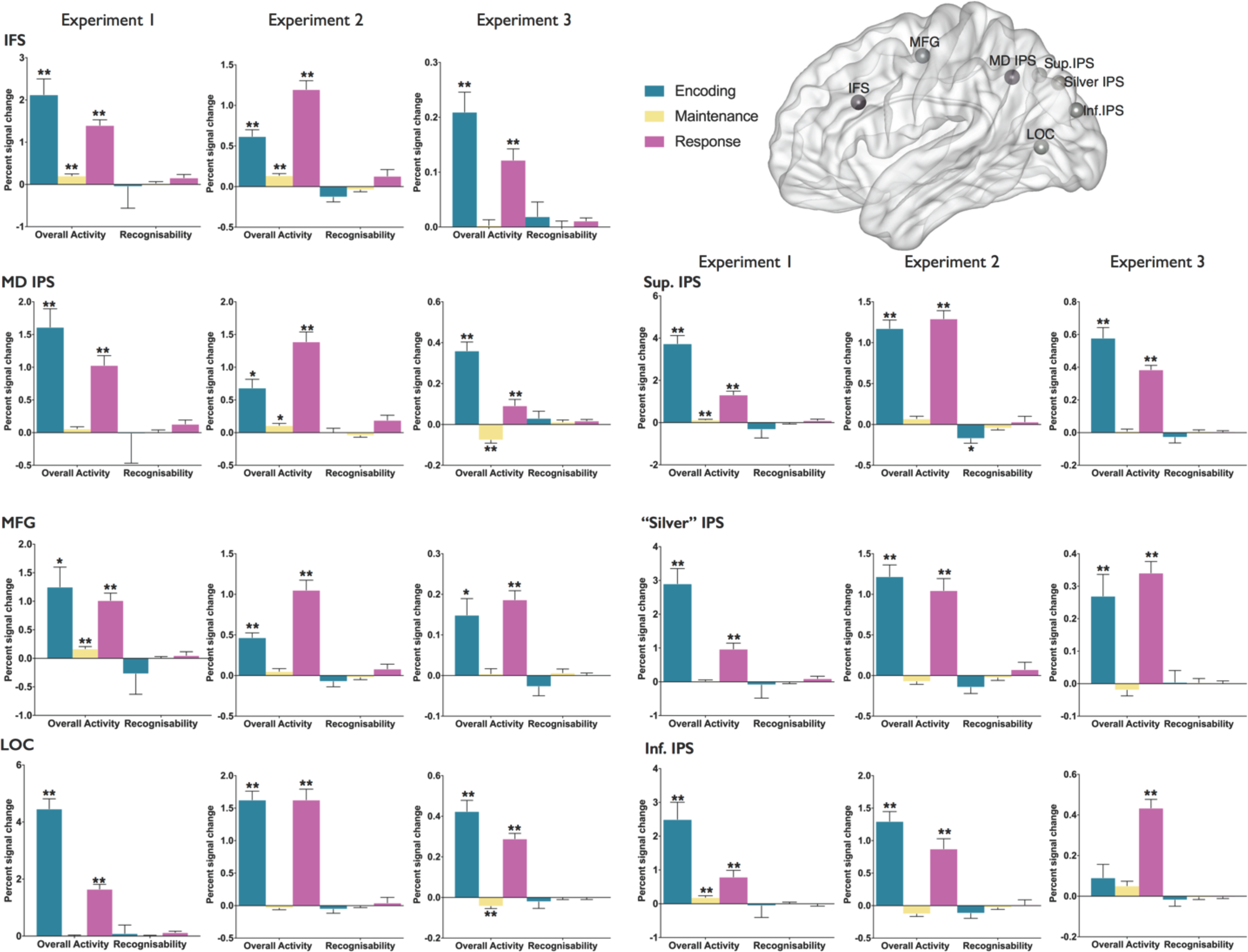
Plots depict mean percent signal change from overall activity (versus implicit baseline), and recognisability contrasts, in the encoding, maintenance and response phases of each experiment. For each ROI, left hand plot represents data from Experiment 1, middle plot from Experiment 2 and right hand plot from Experiment 3. One sample t-test 2-tailed significance level ** p<0.001, * p<0.007 Bonferroni corrected for multiple comparisons across regions. Error bars represent standard error.

**Fig 5.**
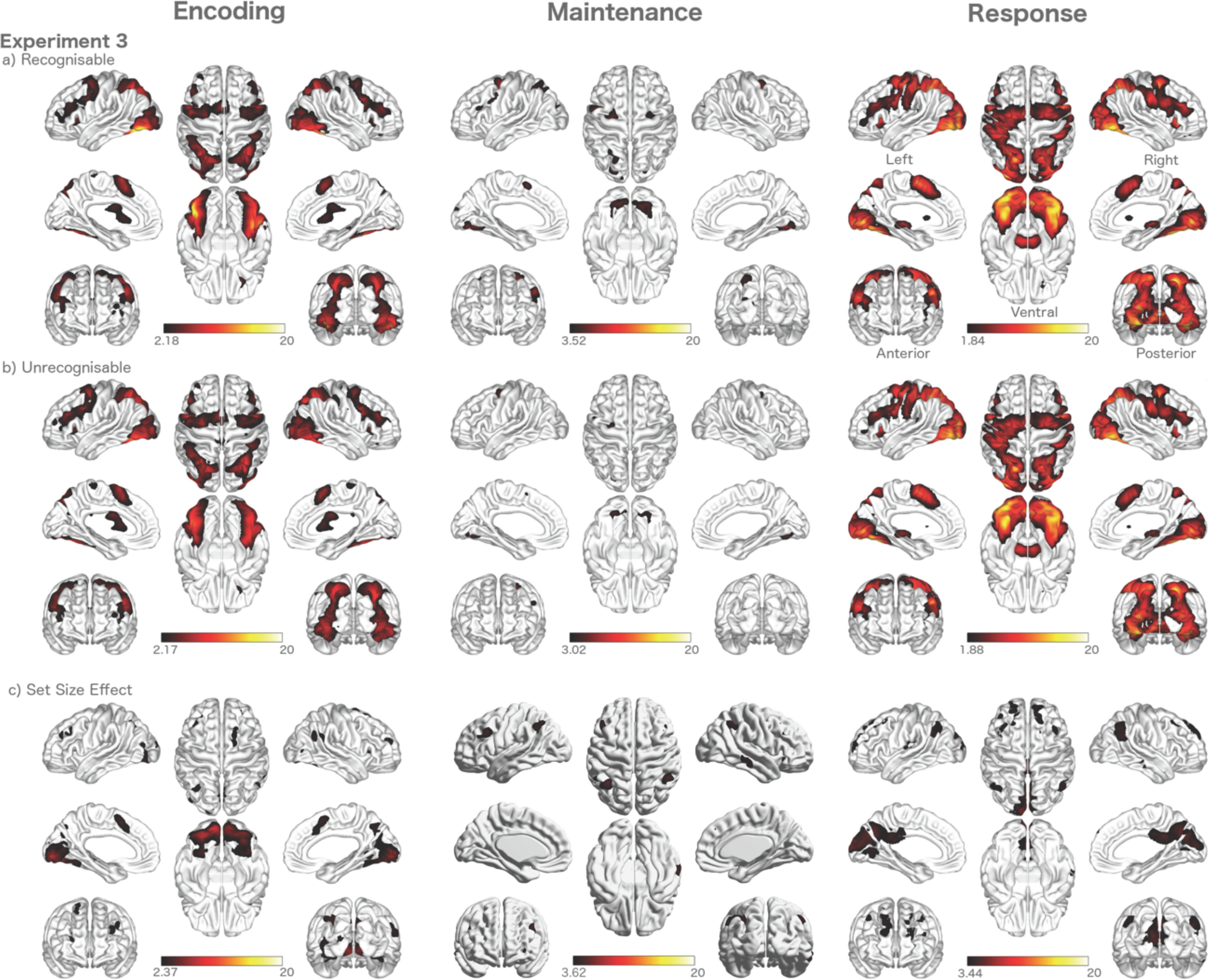
Experiment 3: Activity associated with recognisable objects (row a) and unrecognisable objects (row b) in the encoding, maintenance and response phase of the continuous report task. Significantly greater activity in the encoding, maintenance and response period in set size 2 compared to set size 1 trials. Colour bars represent t-values. FDR <0.05

**Fig 6.**
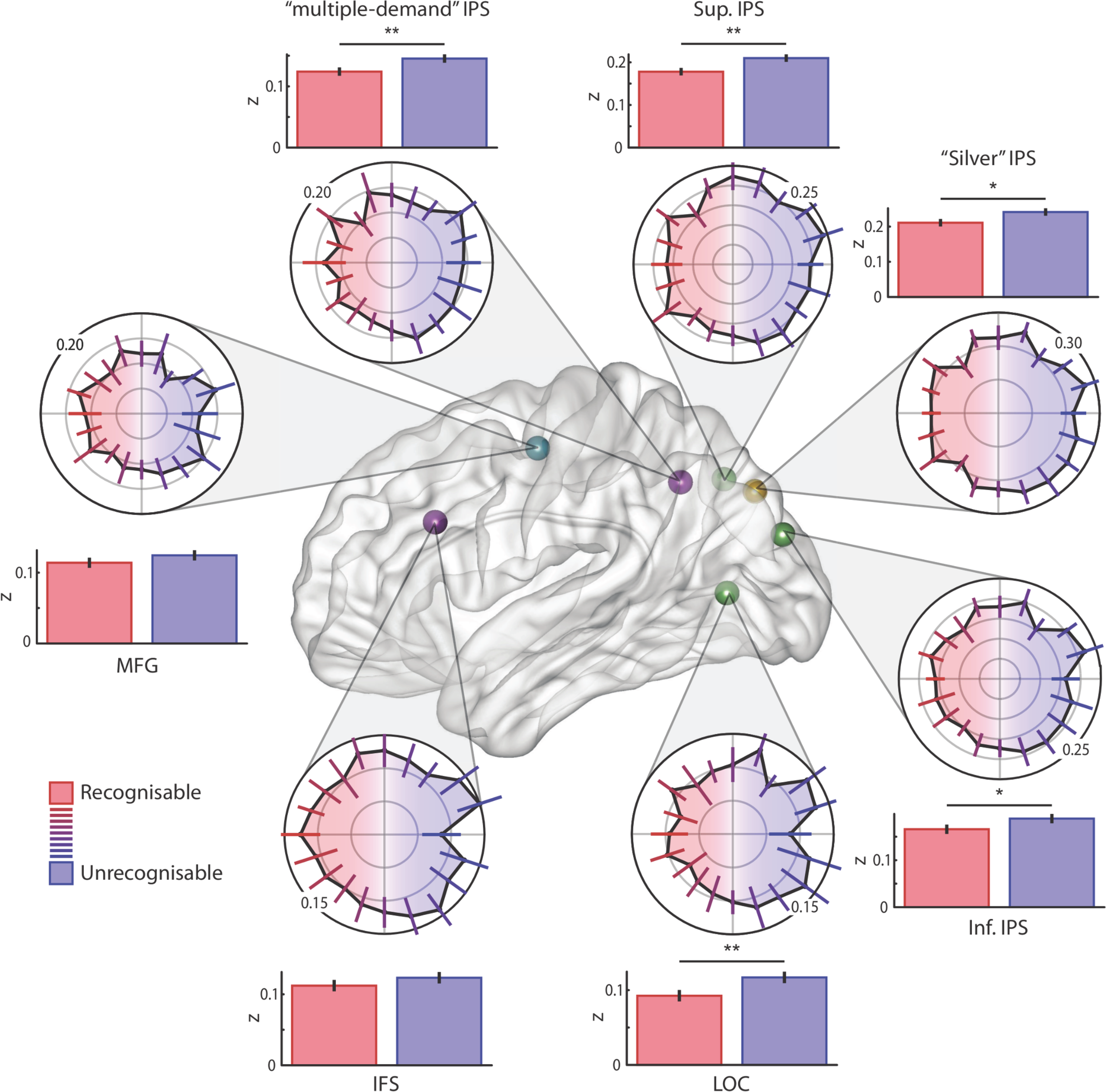
Multivariate representations are more variable for recognisable than unrecognisable objects. Polar plots show mean pattern replicability (Fisher-z-transformed Pearson correlation coefficients) across multiple presentations of an item, separately for each distortion step. Bar plots summarize the mean replicability for more recognisable and less recognisable distortion levels, which are compared by paired t-tests: * indicates p<.05, ** indicates p<.007, surviving Bonferroni correction for multiple comparisons across regions. Error bars are 95% within-subject confidence intervals.

In addition to conventional Student’s t-tests, Bayesian one sample t-tests were conducted in order to assess the likelihood that the data support the null hypothesis of no difference in activity between recognisable and unrecognisable trials. Unlike conventional significance testing, Bayesian one-sample t-tests provide a Bayes factor which is an easily interpretable probability of the alternative or the null hypothesis, based on the observed data. A common rule of thumb for the interpretation of Bayes factors is that a value greater than 3 is considered ‘some evidence’; greater than 10 is considered ‘strong evidence’ and greater than 30, ‘very strong evidence’ in favour of the null or alternative hypothesis (we have reported Bays factors as the natural logarithm of the odds of the alternative hypothesis over the null hypothesis). The interested reader is referred to Rouder et al. (2009) for an excellent primer on Bayesian t-tests. Bayesian t-tests were conducted in JASP statistical software (Love, J., et al. 2015). For all reported Bayesian t-tests, we used the default prior on effect size (Cauchy distribution, centred on zero, with rate r=0.707).

### 1.2 Results

#### 1.2.1 Behavioural Results

Performance was examined with a repeated-measures random-effects design with factors of change condition (CC, WC) and item recognisability (recognisable, unrecognisable). As expected, capacity was significantly greater in the CC than WC trials (*F*(1,17)=135.59, *p*<.001, η^2^_p_=.89). There was also a main effect of recognisability indicating memory capacity was significantly greater for recognisable compared to unrecognisable items (*F*(1,17)=10.58, *p*<.005, η^2^_p_=.38). There was also a significant interaction (*F*(1,17)=4.69, *p*=.045, η^2^_p_=.22) between change condition and item recognisability (Fig 2a). Paired samples t-tests confirmed the interaction was driven by greater capacity for recognisable items compared to unrecognisable items in the WC condition, t(17)=3.41, p=.003, *d*=.80, but not the CC condition, t(17)=.94, p=.36, *d*=-.22.

#### 1.2.2 fMRI Results

##### 1.2.2.1 Whole brain results

Figure 3a-b) shows activity evoked by recognisable and unrecognisable target item trials in the encoding, maintenance and response phases of the task. As is apparent from their similarity, no significant difference was found between recognisable versus unrecognisable stimuli in the encoding or maintenance period, providing no support for the hypothesis that recognisable stimuli recruit additional regions. Only in the response period was there some evidence of a difference, with more activity for recognisable objects in the medial frontal cortex, lateral occipital regions and the fusiform gyrus (Supplementary Fig S1a). No difference in activity was found between WC and CC trials in the encoding period, with limited effects seen in the maintenance and response period (Supplementary Fig S2). There was no interaction with recognisability.

Encoding of complex objects, regardless of recognisability, was associated with a classical fronto-parietal network (Linke et al. 2011) and occipital to ventral stream activity, including the fusiform gyrus. Much of this activity, with the exception of lateral occipital cortex, persisted throughout the maintenance period in the absence of visual stimulation. The response period activated the encoding network and additional regions, reflecting the decision and motor requirements of this phase of the task.

##### 1.2.2.2 ROI results

The ROI analysis (Fig 4) confirmed significant activity in the encoding and response period in all regions in the contrast against implicit baseline. In the maintenance period activity was significant in the inferior and superior IPS and the inferior frontal sulcus and middle frontal gyrus. There were no significant main effects of recognisability (Fig 4) or change condition (Supplementary Fig S3) in any epoch of any ROI in two-tailed one sample t-tests. Bayesian one sample t-tests were conducted in order to measure the likelihood that the data support the null hypothesis of no difference in activity between recognisable and unrecognisable trials. Across all ROIs (Table 1), Bayes factors showed 3-4 times stronger support for the null hypothesis of no difference in the encoding or maintenance period responses for recognisable compared to unrecognisable items. In the response period, only activity in the lateral occipital cortex ROI showed some evidence in support of the alternative hypothesis. For the WC-CC contrasts, with the exception of the superior-IPS at encoding, Bayes factors (Table 1) supported the null over the alternative hypothesis across all epochs and ROIs.

### 1.3 Discussion

The visual features of recognizable items were better remembered than those of unrecognizable items. The results were consistent with the effect being an increase in precision, in that the effect of recognisability was found for small (WC) but not large (CC) changes. This finding mirrors the perceptual expertise advantage for VSTM of faces (Curby and Gauthier 2007; Scolari et al. 2008; Curby et al. 2009; Lorenc et al 2014), but extends it across multiple object categories.

Despite the behavioural benefit for recognizable objects, there was no evidence of recruitment of additional brain regions, or an increase in the strength of activity, for the maintenance of recognizable compared to unrecognizable complex objects. This suggests that the benefit may be due, instead, to more efficient coding within the same regions, but raises the question of whether the experiment had sufficient power to detect a difference in activity. There does appear to be tight replication across the conditions (Fig 3a-b), and the Bayesian t-test provided more support for the null hypothesis than the alternate hypothesis (Table 1). Nonetheless, a stronger confirmation could be to demonstrate that activity can be detected from a manipulation that has a well-established effect, with similar power to the recognizable versus unrecognizable contrast.

In experiment 2, we therefore included an additional manipulation of set size, which is known to affect precision (Bays & Hussain, 2008), and to modulate neural activity (Todd & Marois, 2004; Mitchell & Cusack, 2008). We intermixed CC and WC trials so that the precision required to do the task could not be anticipated. Therefore, any effects on precision should be the result of the stimuli and not strategy encouraged by blocking the trials.

A second extension in experiment 2 was introduced, to probe the possibility that the difference in performance observed was a result of encoding rate varying between recognizable and unrecognizable objects (Eng et al. 2005; Brady et al. 2016). We manipulated encoding time, and so could test whether the visual features of less recognizable objects might be remembered just as well as those of recognizable objects when given longer to encode them.

## 2 Experiment 2

### 2.1 Methods

#### 2.1.1 Participants

Twenty-five new participants (14 male, aged 18-48, mean age 29, SD 7.93) were recruited in the same manner and with the same exclusionary criteria as reported in Experiment 1 (section 1.1.1).

#### 2.1.2 Change Detection Behavioural Paradigm

The sample was presented for 0.5, 1.25 or 2 seconds on a uniform grey background (Fig 1b). On set-size 1 trials, a single item was presented 30mm to the left or right of fixation at a visual angle of 1.81º. On set-size 2 trials, items were presented simultaneously on both sides. Each item was either recognisable or unrecognisable. The sample was followed by a jittered maintenance period (range of 1-9 seconds, randomly distributed) in which participants fixated on the central cross. Participants used a button box to indicate whether a subsequent centrally presented probe item matched any of the items in the sample display. The button-response mapping was counterbalanced across participants. Participants were given a 10 second window in which to make a response before the experiment moved on to the next trial. Half of trials were change trials. In CC trials, the change was in the identity of the stimulus, maintaining the level of recognisability. In the WC trials, the identity of the sample was held and the level of recognisability was changed. Change type, set-size, correct response, encoding duration, and recognisability were counterbalanced and randomly intermixed. Participants completed 168 set size 1 trials and 168 set size 2 trials randomised across four blocks.

VSTM capacity (K) was again calculated using Cowan’s formula as appropriate for single-probed recognition tasks (Rouder et al. 2011). The CC and WC conditions were intermixed and so only distinguishable at the response phase and when a change occurred. The no-change trials were therefore common to both conditions, and capacity estimates for each condition were calculated relative to *all* no-change trials (Awh et al. 2007).

#### 2.1.3 Image acquisition, processing and analysis

Functional imaging data acquisition and preprocessing were as described in Experiment 1. Data from five participants were excluded from the imaging analysis due to excessive movement in the scanner (greater than 5mm translation or 5 degrees rotation), yielding N=20 for the final analysis. These data were included in the behavioural analysis, with the exception of 1 of these subjects whose raw behavioural data were also corrupted. Statistical analysis was the same as Experiment 1 except that regressors of interest modelled recognisable and unrecognisable target item trials; set size 1 and set size 2 trials; short, medium and long encoding duration trials and CC and WC change trials in the three phases of the task (encoding, maintenance and response). The ROI analysis was also as conducted in Experiment 1.

### 2.2 Results

#### 2.2.1 Behavioural Results

##### Repeated measures ANOVA

A four-way repeated measures ANOVA examined the effects of change condition (CC, WC), encoding duration (0.5, 1.25, 2 second encoding), set size (1 or 2 items) and item recognisability (recognisable, unrecognisable) on estimates of VSTM capacity (K) in the change detection task (Fig 2b). There were significant main effects of change condition (F(1,23)=191.59, p<.001, η^2^_p_=.89) and encoding duration (F(2,46)=5.88, p<.01, η^2^_p_=.20). Notably, there were clear main effects for both set size (F(1,23)=46.36, p<.001, η^2^_p_=.67), and recognisability (F(1,23)=199.81, p<.001, η^2^_p_=.90) with large and comparable effect sizes.

The effect of change condition interacted significantly with recognisability (F(1,23)=125.91, p<.001, η^2^_p_=.85) and set size (F(1,23)=62.22, p<.001, η^2^_p_=.73), but not encoding duration (F(2,46)=2.13, p=.131, η^2^_p_=.09). There were significant interactions between recognisability and encoding duration (F(2,46)=3.85, p<.05, η^2^_p_=.14), recognisability and set size (F(1,23)=33.59, p<.001, η^2^_p_=.59) but not encoding duration and set size (F(2,46)=0.16, p=.86, η^2^_p_=.01).

There were significant interactions between change condition, recognisability and encoding duration (F(2,46)=5.17, p<.01, η^2^_p_=.18) and change condition, recognisability and set size (F(1,23)=33.71, p<.001, η^2^_p_=.59).

##### Post-hoc t-tests

In the CC condition, VSTM capacity was significantly greater for recognisable items compared to unrecognisable items whether a single item (t(23)=3.50, p=.002, *d*=-11.90) or two items (t(23)=3.16, p=.004, *d*=-6.95) were presented (Fig 2b i). In the WC condition (Fig 2b ii), capacity was also greater for recognisable compared to unrecognisable items in single (t(23)=16.19, p<.001, *d*=-1.75) and 2 item trials (t(23)=11.51, p<.001, *d*=0.28). The average *difference* in capacity between recognisable and unrecognisable items was smaller in the CC condition than the WC condition for set size 1 (t(23)=-11.87, p<.001, *d*=-2.70) and set size 2 (t(23)=-10.07, p<.001, *d*=-2.26) trials.

In the CC condition (Fig 2b iii) paired samples t-tests confirmed significantly greater capacity for recognisable compared to unrecognisable items when the encoding duration was 0.5 seconds (t(23)=4.22, p<.001, *d*=-5.19) and 1.25 seconds (t(23)=2.43, p=.02, *d*=-5.57) but not 2 seconds (t(23)=1.38, p=.18, *d*=-9.79). Capacity for unrecognisable items increased when the encoding duration was increased from 0.5 seconds to 2 seconds, t(23)=-2.33, p=.003, *d*=-5.59. In the WC condition (Fig 2b iv) capacity was significantly greater for recognisable items compared to unrecognisable items whether the encoding duration was 0.5 seconds (t(23)=11.64, p<.001, *d*=-0.03); 1.25 seconds (t(23)=7.94, p<.001, *d*=-0.82) or 2 seconds (t(23)=9.42, p<.001, *d*=-0.36). Memory performance improved for unrecognisable items when the encoding duration was increased from 0.5 seconds to 1.25 seconds (t(23)=-4.47, p<.001, *d*=-1.42) but not from 1.25 to 2 seconds (t(23)=1.56, p=.13, *d*=-0.76).

#### 2.2.2 fMRI Results

##### 2.2.2.1 Whole brain results

Repeated-measures random effects analysis were used to examine the effect on activity of the factors change condition (CC, WC), encoding duration (0.5, 1.25, 2 second encoding), set size (1 or 2 items) and item recognisability (recognisable, unrecognisable), in the three phases of the task (encoding, maintenance, response). A broad fronto-parietal network was recruited by the encoding of complex objects, irrespective of whether they were recognisable or unrecognisable (Fig 3c-d). This replicates Experiment 1, and mirrors activity associated with encoding of simple objects (Linke et al. 2011). There was also clear fusiform activity in the encoding period, but in contrast to experiment 1 this was not robustly sustained through the maintenance period. Further replicating the results of Experiment 1, there was no detectable difference in activation associated with the encoding or maintenance of recognisable compared to unrecognisable trials. In the response period, there was significantly greater activity for recognisable compared to unrecognisable objects in the precuneus and lateral occipital cortex (Supplementary Fig S1b). In the encoding period, there was some evidence of more activity for unrecognisable compared to recognisable objects in small regions of left frontal and occipital cortex (Supplementary Fig S1c). There was no significant difference between WC or CC trials in any epoch.

We also examined the effect of the set size manipulation in the encoding, maintenance and response phase of the task by contrasting activity that was greater for set size 2 compared to set size 1 trials. In the encoding period there was overall greater activity in set size 2 trials in all the key regions of the fronto-parietal network that frequently show increased activity with increasing memory load in VSTM tasks with simple objects (Fig 3e). Much of this is likely to reflect differences in visually evoked responses between conditions. In the maintenance period, set size effects were significant around the inferior frontal sulcus and within the posterior parietal lobe. Assuming effective separation of the encoding and maintenance phase responses, this is expected to more directly reflect differences in memory load. No significant set size effects were apparent in the response period.

##### 2.2.2.2 ROI results

Replicating the first experiment, there was significant activity across all ROIs in the encoding and response period in overall activity contrasts against implicit baseline (Fig 4; Table 2). In the maintenance period, activity was more limited, in line with the whole brain analysis, with only the IFS and MD-IPS showing significant activity against implicit baseline. There was no significant difference in activity in any ROIs or epochs for WC compared to CC trials (Supplementary Fig S3), as was also seen in the first experiment, with Bayes factors indicating more evidence in support of the null hypothesis across the majority of ROIs (Table 2). There was again minimal evidence of a difference in activation between recognisable and unrecognisable trials in any ROIs or epochs (Fig 4), estimated with Student’s and Bayesian t-tests (Table 2). The only exception was the superior IPS, which showed significantly more activity associated with encoding unrecognisable items compared to recognisable items at encoding (Fig 4).

### 2.3 Discussion

Experiment 2 replicates the key findings of experiment 1, in that recognisable objects were more precisely remembered, but did not activate additional brain regions, or activate the same regions more strongly in the encoding or maintenance period. Experiment 2 extended upon the previous experiment in two ways. First, we show that our neuroimaging paradigm is sensitive to the set-size manipulation during memory maintenance as well as encoding. This is in contrast to the recognisability manipulation, which had a behavioural effect of similar magnitude but had no detectable effect in the whole brain or ROI analyses during encoding or maintenance. Although some of the set-size response surely reflects purely perceptual differences during the encoding phase, differences in the maintenance period are more likely to reflect mnemonic processes, consistent with previous literature (Todd and Marois 2004; Brady et al. 2011, 2016). Second, we show that the benefit of recognizable objects is not due to faster encoding, as even quadrupling the encoding duration (from 0.5 s to 2 s) had only a small effect on performance. The difference in capacity for recognizable compared to unrecognizable items was somewhat reduced by increasing encoding duration, but in the WC condition capacity remained dramatically lower for unrecognisable items at all encoding durations, and there was no benefit to performance by increasing encoding duration from 1.25 to 2 s. This suggests that the time given to encode complex unrecognisable items was not the bottleneck to performance (Eng et al. 2005).

The results from experiments 1 and 2 are consistent with recognisability affecting precision, as fine (WC) discriminations showed a larger difference than coarse (CC) discriminations. However, there remains a possible alternative explanation. In both experiments, performance on the CC condition was high, and so the limited benefit for recognizable objects could reflect a ceiling effect. To address this, in a third experiment, we employed a report paradigm (modelled on Zhang & Luck, 2008) that provides a more nuanced measure of the precision of memory representations.

## 3 Experiment 3

We used our novel parametric stimulus transformation to create a circular response space to estimate the resolution of memory representations for a broad range of complex photographic objects. In continuous report tasks, participants are required to report a feature of a remembered item on a continuous scale. The continuous parameter space is modelled to gain an estimate of the precision of the stored item by quantifying the degree of deviation of the reported feature value from the probed feature value, as well the probability that the item is retained at all (Wilken and Ma 2004; Bays and Husain 2008; Zhang and Luck 2008).

### 3.1 Methods

#### 3.1.1 Participants

Forty-five new participants (21 male, aged 19-59, mean age 30) were recruited in the same manner and with the same exclusionary criteria as reported in Experiment 1 (section 1.1.1). Visual acuity was additionally verified using the Functional Acuity Contrast Test, a sensitive test of functional visual acuity (Ginsburg 2003).

#### 3.1.2 Continuous Report Behavioural Paradigm

The stimulus set was made of 20 steps of diffeomorphic transformation of each original (recognisable) stimulus in a circular parameter space, with the midpoint of the continuum being the most distorted (unrecognisable) image. These 20 steps formed a response wheel for the continuous report paradigm. The most distorted, unrecognisable stimulus lay opposite the intact, recognisable image in the circular response space.

The sample array was presented for 1 s. Items appeared 30 mm to the left or right of fixation subtending a visual angle of 1.81º (Fig 1c). The position and the identity of the sample items were counterbalanced. The level of distortion of the target stimulus was selected from a uniform random distribution centred five steps either side of the original stimulus (recognisable) or the most distorted stimulus (unrecognisable). The sample array was followed by a jittered maintenance period (randomly selected from a range 1-9 s) in which participants fixated on a central crosshair. In the response period, the probed item was indicated by an ‘X’, surrounded by the stimulus response wheel for a maximum of 20 s; a response advanced the trial immediately. The response wheel was presented with a randomly placed square selection box over one of the transformations. The orientation of the response wheel was random on every trial. The selection box could be moved in a clockwise or anti-clockwise direction around the response wheel with the button box until a selection was made with a central button. Holding down the buttons accelerated the movement of the selection box. A jittered inter-trial interval (randomly selected from 0.5-5 s) followed in which participants fixated on the central cross. The task compromised 160 set size one and 80 set size two trials across four blocks. Each item was presented in isolation four times across the experiment to enable multivariate pattern analysis on stimulus repetitions. Set size and recognisability manipulations were counterbalanced and randomly intermixed.

#### 3.1.3 Behavioural analysis

The distribution of errors was fitted with a probabilistic mixture model using code from Paul Bays (http://www.paulbays.com/code/JV10/index.php). This estimates the contribution of target responses, guesses and mislocalisation errors to the overall distribution of response errors (Bays and Husain 2008; Bays et al. 2009). Response errors are calculated as the deviation of the reported response value from the target value, in either direction of the circular response space. These are fit by a von Mises distribution centred on zero error, and a uniform distribution to model guesses. The von Mises distribution is a circular analogue of the normal distribution, that allows modelling of responses from a circular response space. Mislocalisation errors are not relevant in the current experiment since the response wheel only contained response options relevant to the target item. The function returns maximum likelihood estimates of the parameters of the model: an estimate of the concentration parameter, or narrowness, of the von Mises distribution, which describes response precision, and the relative probability of the three types of response. The functions were adapted to improve the estimation of the concentration parameter using a method shown to give a better approximation (Hassan et al. 2012) than the Best and Fisher (1981) method. Prior to modelling, the error distributions were reflected and then averaged to create symmetrical distributions centred on zero error.

The concentration parameter is converted to circular standard deviation (SD), a measure of the width of the distribution and the inverse of the precision of the representation. The probability that the item was not present in memory can be estimated from the height of the uniform distribution and the remaining probability is the probability that the item is retained in memory (Pm). VSTM capacity, measured as the items-worth of information retained (K) can also be estimated from this parameter by simply multiplying Pm by set size (Zhang and Luck 2008; Gold et al. 2010).

#### 3.1.4 Image acquisition, processing and analysis

Functional imaging data acquisition and preprocessing were the same as described in Experiment 1 and 2. Eighteen of the 45 participants were excluded from further analysis due to excessive movement in the scanner (greater than 5 mm translation or 5 degrees rotation), resulting from incorrect positioning of the projection of the stimuli on the screen. Although the MR operator checked that all four corners of the screen were visible, the lower centre was obscured by the curve of the head coil, which caused these 18 participants to move their heads during trials to view the bottom of the response wheel. These subjects are included in the behavioural analyses because they repositioned themselves to see the entire response wheel, enabling them to complete the task. This repositioning caused significant movement artefacts that led to them being excluded from the imaging analysis. Individual subject and group analysis was as described in Experiment 1 and 2.

### 3.2 Results

#### 3.2.1 Behavioural Results

Repeated-measures tests examined the effect of set size, recognisability and their interaction on estimates of precision (SD) and the items worth of information retained (K). There was a significant reduction in SD for set size 1 compared to set size 2 trials (Table 3, Fig 2c i) indicating a single item was stored with greater precision than two items. K increased significantly from set size 1 to set size 2 in line with typical set size effects seen in VSTM experiments. In contrast, in recognisable, compared to unrecognisable item trials, there was no significant difference in K. Recognisable items were remembered with significantly greater precision than unrecognisable items (Table 3, Fig 2c ii). There was no significant interaction between the effects of recognisability and set size on SD (F(1,39)=.35, p=.56, η^2^_p_=.01) or K (F(1,39)=1.05, p=.31 η^2^_p_=.03).

**Table 3.** Paired samples t-test statistics between estimates of memory performance in set size 1 compared set size 2 trials (left columns) and recognisable compared to unrecognisable items (right columns).

|  | Set Size |  |  | Recognisability |  |  |
| --- | --- | --- | --- | --- | --- | --- |
|  | t | p-value | Cohen's d | t | p-value | Cohen's d |
| **SD** | -3.25 | .002 | -1.90 | -5.17 | .001 | -2.30 |
| **K** | -9.54 | .001 | -2.62 | -1.01 | 0.32 | -1.45 |

#### 3.2.2 fMRI Results

##### 3.2.2.1 Whole brain results

Activity in the encoding, maintenance and response period of this continuous report paradigm was strikingly similar in spatial distribution to the same contrasts in the two change detection tasks (Fig 5a-b). The pattern of fronto-parietal activity in the encoding period also matched that seen for encoding of simple objects in VSTM (Linke et al. 2011). Replicating experiment 1, fusiform activity in the encoding period was sustained through the maintenance period in the absence of visual stimulation. This region also showed set size effects, with increased activity for set size 2 trials compared to single item trials (Fig 5c). Set size effects were also seen in the maintenance period in similar regions to that seen in the same contrast in Experiment 2, including the posterior parietal lobe. The response period showed a similar, although more widespread, pattern of activation to the encoding period, involving the frontoparietal network and ventral temporal regions. As in experiments 1 and 2, there was not significantly greater activation for recognisable compared to unrecognisable items in the encoding and maintenance period, extending to the response period in this experiment. There was also no significantly greater activity for unrecognisable compared to recognisable items in the encoding, maintenance or response periods.

##### 3.2.2.2 ROI results

There was significant overall activity in all ROIs in the encoding and response period, with the exception of the inferior IPS at encoding (Fig 4). This replicates both change detection experiments. In the maintenance period there was a significant suppression of activity in the LOC and MD-IPS. In support of the two change detection experiments, there was again minimal evidence of a difference in activation between recognisable and unrecognisable trials in any ROIs or epochs (Fig 4), estimated with Student’s and Bayesian t-tests. The only exception was ‘MD IPS’ during the response phase, for which the Bayesian test showed some evidence for more activity associated with recognisable items compared to unrecognisable items. During the encoding and maintenance phases, Bayes factors indicated 2.3-4.9 times more support in favour of the null hypothesis than the alternative hypothesis. This was in contrast to set size effects which were evident in the inferior and MD IPS in the encoding period and the IFS and MD-IPS during maintenance (Supplementary Fig S3). In the maintenance period there was also significantly more activity for single item trials compared to two item trials in the inferior IPS and LOC.

### 3.3 Discussion

The results replicate the two change detection experiments and generalise to a continuous report paradigm. Object recognisability affected the precision (SD) but not the number of distinct items (K) that could be retained. Importantly, here we used a more direct measure of precision and confirmed that the behavioural advantage of recognition to memory performance is in the precision of memory representations. There was no evidence of ceiling effects, supporting the interpretations drawn in the change detection experiments. Despite this substantial behavioural advantage, there was no clear evidence of recognisable objects being associated with increased (or reduced) brain activation, compared to unrecognisable objects, nor recruiting additional brain regions. In contrast, and replicating experiment 2, the behavioural set size effects were accompanied by increased neural activation when 2 items were encoded in comparison to a single item.

Across all three experiments there was a clear behavioural advantage of recognisability, but with no consistent difference in the neural activity associated with the recognisable compared to unrecognisable items. The set size effects confirmed that the neuroimaging data were sensitive enough to detect behavioural effects of comparable size. Therefore, we found no evidence in support of the hypothesis that VSTM for recognisable items recruits additional brain regions, or activates regions more strongly. In the final experiment, we went on to test whether there was information in the fine-scale patterns of activity associated with recognisable compared to unrecognisable objects, that would not be seen at a univariate level, but may help to explain the clear, replicated behavioural pattern we observed. Multivariate methods have the sensitivity to detect representational content, contained within patterns of brain activity, that is not available to standard univariate methods (Mur et al. 2009). These methods have been usefully applied to decode the contents of VSTM e.g. (Ester et al. 2009; Harrison and Tong 2009; Serences et al. 2009; Emrich et al. 2013; Bettencourt and Xu 2015). In the context of episodic memory, representational similarity analysis (RSA, Kriegeskorte, Mur, and Bandettini 2008) has revealed greater pattern similarity between repetitions of a subsequently remembered stimulus compared to a forgotten stimulus (Xue et al. 2010).

## 4 Representation Similarity Analysis

To test the hypothesis that better VSTM for recognisable objects is supported by quantitatively different patterns of activity compared to unrecognisable objects we used RSA to measure pattern similarity between repetitions of the same stimuli within the ROIs probed with univariate analysis in the previous three experiments.

It is increasingly recognised that brain signal variability may be an important indicator of neural processing efficiency (McIntosh et al. 2008; Garrett et al. 2011). In brain development and aging, brain signal variability, whether measured with EEG or fMRI, correlates with behavioural performance (McIntosh et al. 2008; Garrett et al. 2011). Greater signal variability is associated with more consistent reaction times and more accurate behavioural performance (McIntosh et al. 2008). This may reflect a wider range of functional brain states available to support behavioural performance. In the context of memory, the levels-of-processing model posits that stimuli that evoke rich contextual associations are remembered better (Craik and Lockhart, 1972). These contextual associations tend to vary from trial to trial, as encapsulated in the encoding variability principle (Bower et al. 1975). Applying this theoretical framework to our experiment, lower pattern similarity would be expected between repetitions of the recognisable stimuli as they will evoke richer but more variable contextual associations, than unrecognisable ones. However, in a study of the lag effect on episodic memory, Xue et al., (2010) found that stimuli are better remembered if they evoke more consistent activation patterns across repeated presentations. According to this perspective, the recognisable stimuli might evoke higher pattern similarity than the unrecognisable ones, as they are better remembered.

### 4.1 Image processing and analysis

Data from Experiment 3 were remodelled and RSA performed. Preprocessing was as described in Experiment 3 except that images were not smoothed. Each phase of each trial was now modelled separately. As before, events were convolved with the canonical hemodynamic response function. Motion parameters were included in the model. A high pass filter with a cut-off of 128 s was applied. For each event, a t-test compared the regression coefficient against the implicit baseline. Unfortunately, the amount of temporal spacing between events was not sufficient to reliably estimate betas for single epochs of single trials (see e.g. Abdulrahman & Henson, 2016), therefore, to avoid instability in fitting individual events, t-maps from the encoding, maintenance and response phases were averaged per trial, and the resultant t-maps used as the input to the RSA (Misaki et al. 2010).

We conducted the RSA analysis on the ROIs defined and analysed in the first 3 experiments. For each ROI, a correlation matrix compared multivoxel patterns between all pairs of trials on which a single item was presented. Pearson correlation coefficients were Fisher-z-transformed, and then all six pairwise correlations amongst the four repeats of a given stimulus at a given distortion level were averaged. Finally, this measure of pattern replicability was averaged across different stimulus identities. Paired t-tests across subjects were used to compare pattern replicability between recognisable and unrecognisable stimuli. Finally, we conducted correlations across subjects, to relate behavioural performance (P_m_ and SD) with mean pattern replicability across ROIs, separately for recognisable and unrecognisable objects.

### 4.2 Results

Across all ROIs, the trend was for pattern replicability to be lower for more recognisable than for less recognisable objects. This is illustrated by the polar plots in figure 6, which show pattern replicability at each distortion level, and by the bar plots which summarise these values across the most recognisable and least recognisable levels. The difference is not significant in either of the two frontal ROIs, but is significant in all of the occipital and parietal ROIs (“multiple-demand” IPS: t(26)=3.20; Sup. IPS: t(26)=3.69; “Silver” IPS: t(26)=2.92; Inf. IPS: t(26)=2.33; LOC: t(26)=3.28; all p<0.05, two-tailed, uncorrected for multiple comparisons) and is especially robust in the superior IPS and LOC (p<0.007, surviving Bonferroni correction for multiple comparisons).

We correlated behavioural measures of VSTM performance (P_m_ and SD) for recognisable and unrecognisable objects across subjects with pattern replicability for recognisable and unrecognisable objects estimated from the RSA analysis. This served to relate the measure of neural response replicability to memory performance in congruent conditions. The probability of remembering unrecognisable items (P_m_) was significantly positively correlated with the mean pattern replicability of unrecognisable items across ROIs, r=0.65, p<.001 (Fig 7).

**Fig 7.**
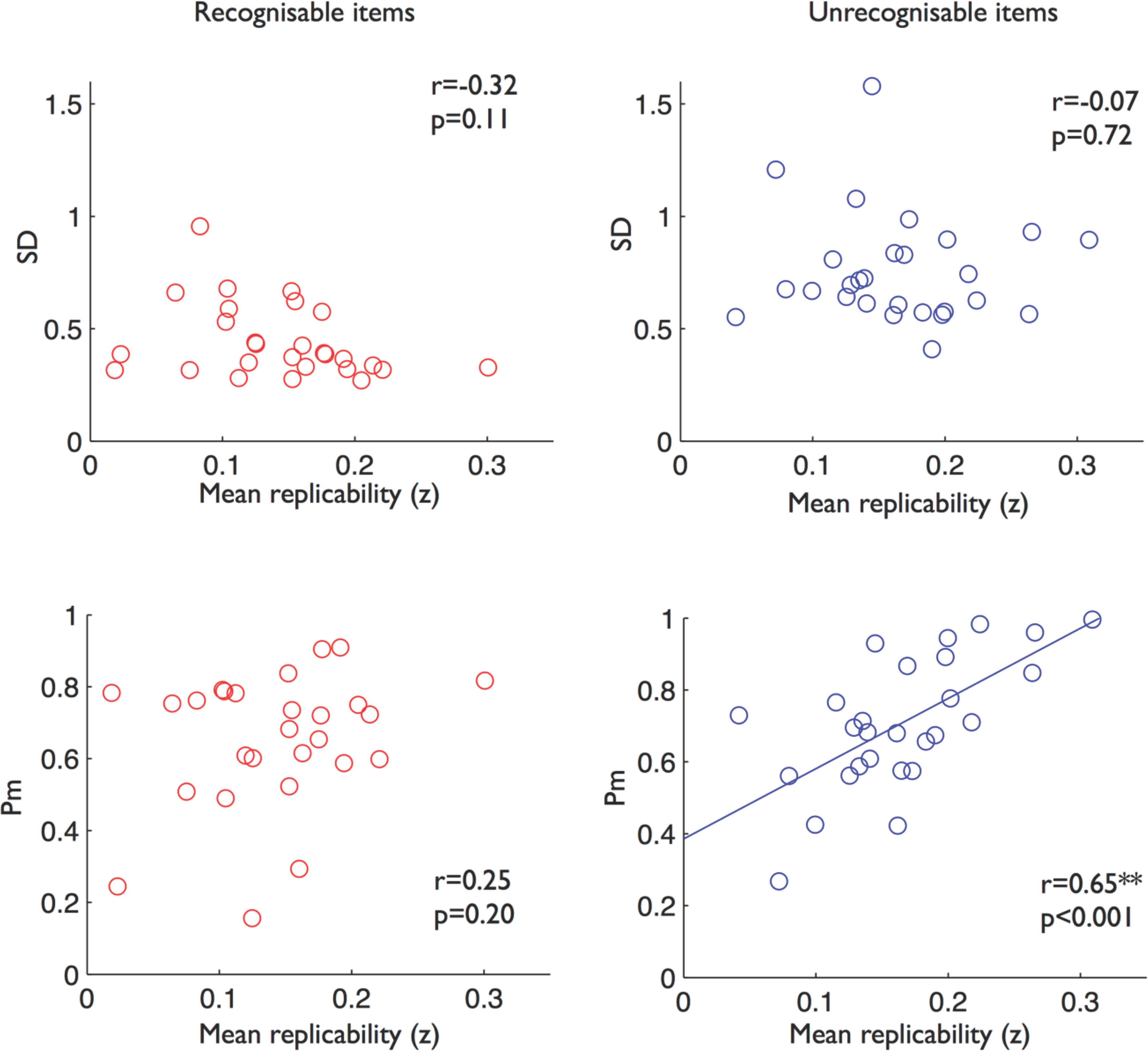
Relationship between measures of memory performance and pattern replicability. Precision (SD) displayed in upper panels; capacity (Pm) displayed in lower panels for recognisable (left panels) and unrecognisable (right panels) items. Performance measures are plotted against mean Fisher-z-transformed Pearson correlation coefficients across all regions of interest. ** indicates p<.0125, surviving Bonferroni correction for multiple comparisons.

### 4.3 Discussion

The RSA analysis found that an image’s recognisability affected neural coding in regions of the intraparietal sulcus and lateral occipital cortex that have been associated with encoding and maintenance of simple objects in VSTM (Todd and Marois 2004; Xu and Chun 2006; Linke et al. 2011). This stands in contrast to the lack of effect of recognisability on overall activity, observed in the univariate ROI analyses of experiments 1-3. The direction of the RSA effect, with greater variability in the patterns evoked by recognisable objects, suggests that this is a result of a richer range of representations associated with objects for which there is a semantic framework (Bower et al. 1975; Garrett et al. 2011). This difference in coding could be responsible for the difference in behavioural performance that was not reflected in any differences in the number or strength of regions recruited.

In addition, we correlated pattern replicability with memory performance for recognisable and unrecognisable objects. In line with Xue et al., (2010), we found greater pattern replicability was associated with higher probability of recalling an item, as measured by Pm. This demonstrates the behavioural relevance of the measure of pattern replicability, and was limited to unrecognisable objects only.

Greater pattern variability for recognisable objects was most robust in the superior IPS and LOC regions, which are thought to reflect the amount of visual detail represented in short-term memory (Xu and Chun 2006). Our results are consistent with broader evidence that multivariate methods have sensitivity to discriminate neural patterns even in the absence of a detectable univariate response (Mur et al. 2009; Serences et al. 2009).

## 5 General Discussion

We used a novel stimulus transformation method to parametrically distort complex photographic images, removing recognisability, without changing perceptual complexity (Stojanoski and Cusack 2014). We then adapted two of the most common VSTM paradigms, developed for simple stimuli, to investigate how complex recognisable objects are maintained in VSTM and their neural correlates.

### Greater precision for recognisable complex objects in VSTM

In both change detection experiments we found a greater effect of recognisability on VSTM performance when fine changes were to be detected, requiring precise memory representations. This is in line with recent findings suggesting that complex, recognisable objects can be stored in VSTM with finer detail compared to simple objects (Brady et al. 2016), extending it to show that, when objects are perceptually matched, recognisability is associated with higher memory precision. This also concurs with prior change detection findings for faces (Curby and Gauthier 2007; Scolari et al. 2008), hypothesised to reflect a wider representational space for upright faces, for which we have perceptual expertise, compared to inverted faces. Here we generalise this effect to a broader range of stimuli. We also extend it to another measure of precision, obtained from modelling the distribution of responses in a continuous report paradigm. This confirmed that recognisable objects are remembered with greater precision but with no advantage in the number of items that can be retained, similar to an advantage seen for upright faces (Lorenc et al. 2014). This advantage in precision may be explained by a wider representational space for recognisable objects (Scolari et al. 2008; Brady et al. 2016).This raised the question as to the neural mechanisms that support VSTM for complex objects and that underlie a behavioural precision advantage for recognisability.

### No consistent evidence for stronger activity or additional regions recruited for memory of recognisable objects

The fronto-parietal network, along with occipital cortex, was found to be central to the encoding of complex objects into VSTM, similar to activity seen in the encoding period in a meta-analysis of VSTM tasks using simple objects (Linke et al. 2011). Activity in frontoparietal regions, and the fusiform gyrus in two of the three experiments, was sustained through the maintenance period. In two experiments set-size was manipulated and several of these regions showed increased activity when two items were presented compared to a single item during the encoding and maintenance periods.

Most strikingly, and consistently across all three experiments, we found no additional activity or recruitment of additional brain regions for the encoding or maintenance of recognisable compared to unrecognisable objects in VSTM, despite the very substantial effect on behavioural performance. Bayesian t-tests consistently confirmed more evidence in favour of the null hypothesis of no difference between univariate responses to recognisable and unrecognisable objects.

We originally hypothesised an increase in activity or the recruitment of additional regions for the encoding or maintenance of recognisable items, but it is also possible that there may be a decrease in activity if recognisable items are more efficiently encoded. In the priming literature, for example, better behavioural performance is seen for repeated or highly familiar items, and it is associated with a decrease in neural activity, termed repetition suppression, thought to reflect more efficient neural processing (Henson and Rugg 2003; Gotts et al. 2012). This did not appear to be the case in the current experiments, with no consistent evidence for reduced activity or a reduced spread of activation for the recognisable compared to unrecognisable objects. Only in experiment 2 were there any voxels with more activity for unrecognisable objects, but they were only in the encoding period, were highly restricted, and inconsistent across experiments.

### More variable neural representations for recognisable objects

Rather than a change in the univariate strength of activity, or recruitment of additional regions, we propose that a richer range of associated semantic representations may support more precise visual memory for recognisable objects. Greater variability in the neural pattern of responses to recognisable objects was seen within the same parietal and occipital regions that are involved in short-term memory of simple or complex objects lacking semantic associations.

Brady et al. (2016) demonstrated that greater VSTM capacity for complex objects was not explained solely by the use of long-term episodic memory, as is commonly assumed. Rather, precise memory of complex objects was facilitated through active VSTM maintenance, as indexed by the electrophysiological marker, contralateral delay activity (Vogel and Machizawa 2004; Ikkai et al. 2010). Contralateral delay activity localises to the posterior parietal cortex (Mitchell and Cusack 2011), where recognisability modulates pattern replicability in the current study.

Our results support the theoretical framework that recognisable objects evoke richer and more variable contextual associations. Variability of brain signals, measured with EEG and fMRI are associated with more accurate and more stable behavioural performance across a range of cognitive tasks, including VSTM tasks (Garrett et al. 2011). This is seen in brain development and in brain aging (McIntosh et al. 2008; Garrett et al. 2011). Brain signal variability increases with brain maturation and is associated with decreased behavioural performance variability, as measured by trial to trial reaction time variability and accuracy (McIntosh et al. 2008). Brain signal variability is a strong predictor of age and a marker of age-related neural inefficiency (Garrett et al. 2011). It is hypothesised that brain signal variability reflects a greater repertoire of brain states which can be easily transitioned between, and which represent optimal neural efficiency (McIntosh et al. 2008). When variability is low, there is less adaptability to uncertainty in the environment. Our results are in line with this interpretation, suggesting recognisability is associated with a wider range of available brain states to support precise VSTM. However, when relating behavioural precision to individual differences in pattern replicability across subjects, we did not see a significant relationship. This cautions that the within-subject association between recognisability and pattern replicability (Fig 6) may not be important in explaining between-subject performance in memory precision, and this would not necessarily be expected (Kievit et al. 2013).

Our finding of greater pattern variability for recognisable items that were better remembered appears somewhat at odds with Xue et al.’s, (2010) study of episodic memory, which concluded that better episodic memory performance was associated with lower variability in response patterns between item repetitions. However, when we tested the relationship between individual differences in pattern replicability and memory performance we found a similar relationship to that demonstrated by Xue et al., (2010). The probability of recalling an item, as indexed by Pm, was positively correlated with pattern replicability for unrecognisable objects. This shows that the measure of pattern replicability has behavioural relevance in terms of differences across individuals, and is in line with the within-subjects across-items association described by Xue et al., (2010). Although we found that memory performance (Pm) was positively correlated with pattern replicability, the reverse relationship might have been expected, ie as pattern replicability decreased (resembling the representational variability of recognisable objects) memory performance would improve. This raises two important points. Firstly, it suggests that memory performance is dissociable in terms of how pattern replicability relates to precision (within subjects) and to the probability of recalling an item (between subjects). This is consistent with several other results in which memory capacity and memory precision have been shown to have independent relationships with intelligence (Fukuda et al. 2010), different trajectories in aging and mental health disorders (Gold et al. 2010) and different neural correlates (Xu and Chun 2006). Secondly, this raises the question as to whether the representation of recognisable and unrecognisable objects might be qualitatively different, such that even when pattern replicability is on the low end for unrecognisable objects, memory performance does not resemble that seen for recognisable objects, as it might if they existed on a continuum. Although our stimuli are well designed to address this question, the current experiments were not. We collapsed across levels of distortion and classified items as either recognisable or unrecognisable. Examining memory performance and pattern replicability for the intermediate levels of distortion might reveal a continuous relationship. Unfortunately we did not have enough repetitions of individual distortion levels to investigate this.

Although we show some convergence with the results from Xue et al. (2010), there are a number of important differences between the paradigms (recall of fine visual detail versus recognition/recall of item identity), and the type of memory tested (memory across seconds versus hours). Furthermore, our experimental manipulation of item recognisability is distinct from the comparison of items that happened to be recalled or forgotten despite being of comparable semantic richness at encoding. Relating these studies would be a valuable direction for future research. Finally, in the RSA analysis we were unable to separate the encoding, maintenance and response epochs within a single trial, so it remains possible that different patterns might be observed in these distinct cognitive stages.

One further limitation of the current work is that multivariate pattern replicability was only measured in the specific context of VSTM for fine visual details. Given the evidence that multivariate coding during VSTM is affected by task (Vicente-Grabovetsky et al, 2012), care should be taken before generalizing these conclusions to other kinds of VSTM task.

## 6. Conclusions

Across three fMRI experiments we demonstrate greater precision of VSTM for recognisable, compared to unrecognisable, complex objects. This improvement in memory performance did not appear to be the result of stronger (or reduced) activity in key brain regions associated with VSTM, or the recruitment of additional brain regions. Rather, recognisable objects evoked more variable neural codes in the intraparietal sulcus and lateral occipital cortex compared to unrecognisable objects, likely reflecting the wider representational space available to recognisable, semantically loaded complex objects in VSTM.

## Funding

This work was supported by the Medical Research Council (grant number MC_U105592690).

